# Brain Connectivity Modelling Through Joint Estimation of Parcels and Gradients

**DOI:** 10.64898/2026.06.23.734045

**Authors:** Aref Miri Rekavandi, Saad Jbabdi, Stephen M. Smith

**Affiliations:** Oxford Centre for Integrative Neuroimaging, FMRIB, Nuffield Department of Clinical Neurosciences, University of Oxford, United Kingdom

**Keywords:** Brain organisation, principal gradients, local and global embedding, resting state fMRI

## Abstract

This paper presents a framework for modelling the topography of whole-brain connectivity in resting-state functional MRI. The aim is to disentangle functional segregation, which manifests as abrupt changes in connectivity, from so-called gradients, i.e., smooth variations in connectivity across the brain. Our core assumption is that functional segregation leads to low-rank structure in the dense (point-to-point) connectome, whereas connectivity gradients imply a sparse and non-low-rank structure in the dense connectome. Our method thus decomposes the connectome into low-rank and sparse components, enabling the integration of local-nonlinear and global-linear embedding strategies. We show that this hybrid model approximates the empirical dense connectome more effectively than purely low-rank or purely gradient approaches. We also find that connectivity gradients derived from this model exhibit strong correspondence with task-based topographic maps. We hope that this approach can provide insight into the organisational principles of brain regions where gradients remain poorly characterised.

## 1 Introduction

Finding the right level of abstraction is a ubiquitous challenge in science. In neuroimaging, modelling brain connectivity at the correct level of granularity remains an unsolved problem. Parcellation-based approaches, where the nodes of the connectome are brain regions, assume that connections within a region are homogeneous [26, 8]. Mathematically, this means that the dense (i.e., point-to-point) brain connectome has a low-rank structure. Linear modelling techniques such as Independent Component Analysis (ICA) can be applied to reveal this low-rank structure [1]. These models are advantageous as they reduce the complexity of the connectome and enable simpler and faster downstream analyses and interpretations. However, it is well known that brain connectivity can spatially vary in a continuous or topographic manner [14, 21]. For example, connectivity profiles within primary areas such as M1 or V1 have 1D or 2D continuous variations that map onto locations on the body or retina. These connectivity profiles cannot be represented by a low-rank structure. A linear decomposition such as ICA would require many components to approximate the spatial variations of connections, while failing to capture the nature and underlying structure of the Dense Connectome (DC). Alternative models have been proposed for modelling these so-called connectivity “gradients”. These models are ususally nonlinear and local, and are often (sensibly) applied within a region of interest [10]. Examples of nonlinear embedding approaches used in gradient methods include Spectral Eigenmaps (SE) [2] and Diffusion Embedding (DE) [5]. These methods have also been applied globally across the cortex to reveal whole-brain connectivity gradients [15], also known as “principal gradients”. However, these whole-brain gradients show great similarity to components obtained through low-rank decomposition (see Figure 1). This raises the question of whether a whole-brain gradient estimation, in its simplest form, truly adheres to its definition and offers novel insights into brain organisation beyond those already captured by low-rank linear analyses.

**Figure 1.**
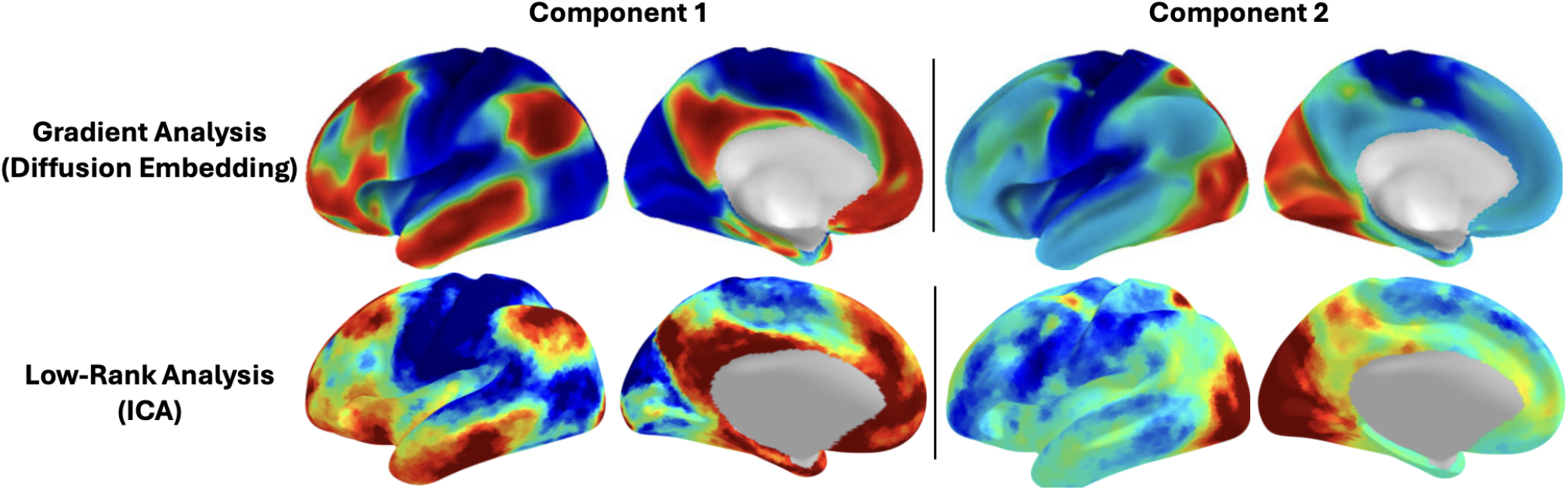
Global principal gradients derived from applying diffusion embedding on 100 randomly selected HCP subjects’ resting-state fMRI data are similar to ICA maps.

In this paper, we discuss the underlying reasons for the observation that local gradient analyses can capture true aspects of brain organisation, whereas global gradients largely replicate patterns already revealed by parcellation-based approaches. To address this, we then propose a novel method, DLS (Dense to Low-rank + Sparse connectomes), designed to uncover genuine (global or local) gradients of brain connectivity separately from the low-rank patterns observed in empirical DCs.

## 2 Preliminaries

In this section, we briefly describe some of the most popular embedding techniques, including linear vs nonlinear and local vs global. Linear models represent the dense connectome via a linear transformation (i.e., a matrix multiplication). The distinction between global and local methods pertains to the nature of the objective function. A global objective aims to transform the DC while preserving its global structure (e.g., covariance) at the expense of local pairwise relationships between voxels. Local methods put more weight on preserving pairwise similarities at the expense of global patterns in the DC.

Throughout this paper, we are given a data matrix denoted by 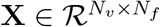 that represents for example a resting-state functional MRI (rfMRI) scan with *N*_*v*_ voxels/vertices and *N*_*f*_ measurements (timepoints) recorded for each voxel/vertex. Embedding techniques applied to such a dataset aim to find a *k*dimensional representation of the data where *k < N*_*f*_ . From this point onwards, to differentiate between the number of low-rank components and the number of gradients in the data, we denote them by *r* and *g* (*r* + *g* = *k*). Often the main goal of applying these embeddings in the context of gradient analysis is to preserve the dominant patterns in the DC. Here the DC matrix **Σ** is the pairwise correlation between all voxels/vertices in the dataset **X**.

### 2.1 ICA (Linear, Global)

The aim of ICA [13] is to find statistically independent components that are a linear transformation of the input data. In fMRI, spatial ICA is usually applied to the time series data **X**. Here we instead (but equivalently) apply ICA to the dense connectome, as we are looking for connectivity-dependent components:

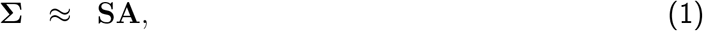

where 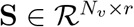 is the sources matrix and consists of spatially independent components and 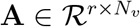 is the mixing matrix. In general, in the context of brain analysis, the independence constraint used in ICA is a more biologically meaningful constraint than the orthogonality constraint used in PCA and its variants [17]. In the case of large DCs, we first apply incremental PCA [18] to compress 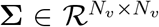 into 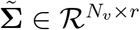 to speed up the process of finding independent components. The estimated ICA maps are a global linear transformation of the input data. We use FastICA [12] for the implementation of ICA.

### 2.2 Gradient Modelling via Nonlinear Embedding

Nonlinear embedding techniques attempt to find a low-dimensional space that best represents a graph (here, the graph is the dense connectome). The objective is for pairwise distances within the low-dimensional space to be a good approximation to the pairwise distances in the original graph. Various embedding methods differ in the way the graph is pre-processed (e.g., hard- or soft-thresholded), and in the loss function that is being optimised. Here we use two popular methods: ISOMAP [20] and UMAP [16].

ISOMAP starts by calculating pairwise “geodesic” distances where the geodesics must traverse the graph through its neighbourhood structure. To compute geodesic distances, a graph is first constructed from the data points, where each point is connected only to its nearest neighbours (i.e., those with minimal pairwise distances). The geodesic distance between two given points is then defined as the distance of the shortest path between them within this graph, traversing only through the edges that connect neighbouring points (confirming that ISOMAP is a global approach, as it cares about all the distances). The algorithm then finds a low-dimensional representation such that Euclidean distances in this representation best approximate the geodesic distances on the graph. This method is thus nonlinear and global.

UMAP also starts by finding each point’s nearest neighbours on the graph. It then converts connection distances to probabilities, where larger distances correspond to lower probabilities, including a local adjustment of these probabilities to improve robustness to outliers. The loss function for finding a low-dimensional embedding then encourages the high- and low-dimensional connection probabilities to be similar in an iterative approach. This makes UMAP nonlinear and both global and local.

Both ISOMAP and UMAP result in a low-dimensional space where voxels with similar connectivity profiles lie close together and dissimilar voxels are placed farther apart. This aligns well with the objective of a gradient analysis, which seeks principal directions of smooth changes in connectivity.

#### Algorithm 1

Dense connectome decomposition into **L** and **S**

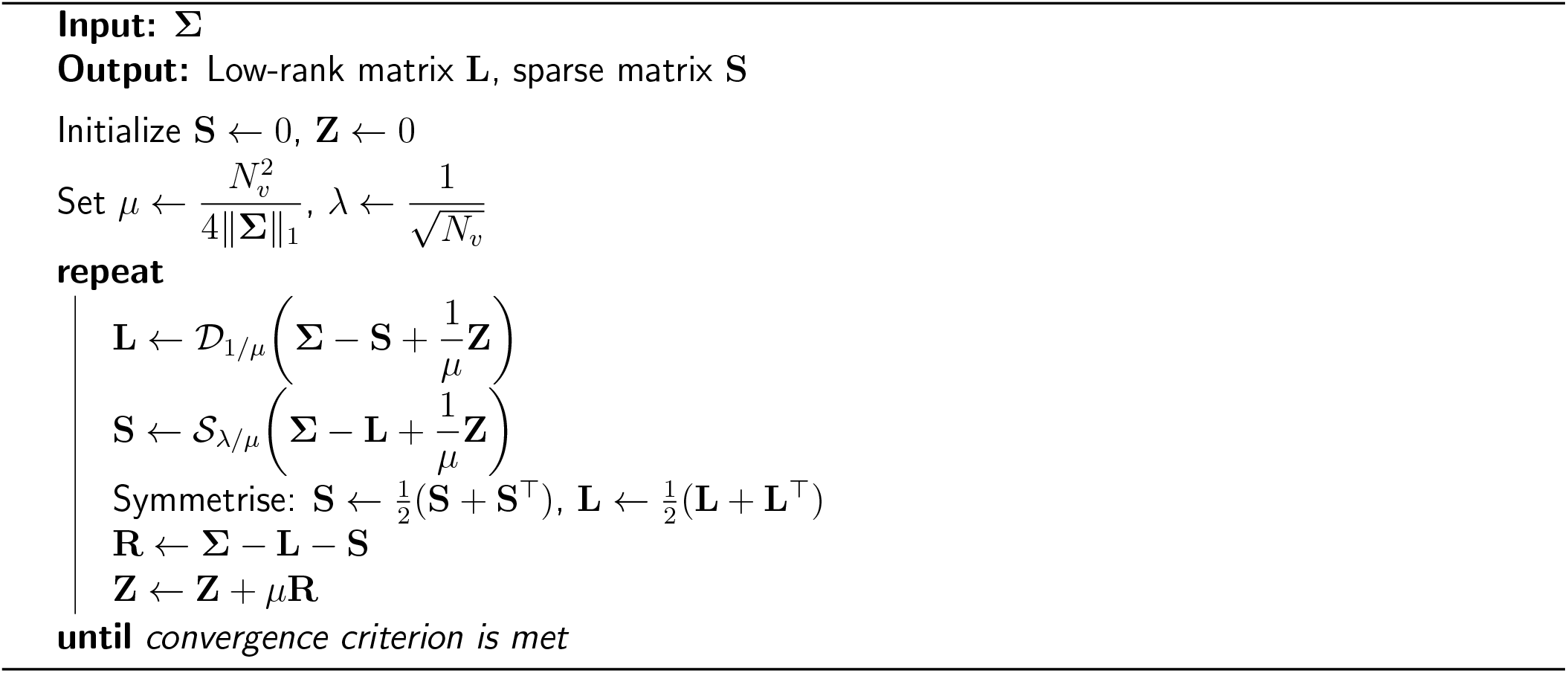

## 3 Methods

In order to disantangle global-linear, low-rank patterns in the DC from connectivity gradients, we propose to use a “low-rank + sparse” decomposition [3]. We then apply nonlinear embedding to the estimated sparse matrix to find the global brain connectivity gradients. Before going into details of the method, we give a brief oveview of the dataset that we used for evaluation.

### 3.1 Dataset

We used publicly available resting-state functional MRI (rfMRI) data from 100 randomly-chosen subjects from the HCP dataset [23]. The full acquisition details and preprocessing pipeline for this dataset are described in [22]. In summary, 3T whole-brain fMRI data were acquired with a spatial resolution of 2 × 2 × 2 mm^3^ and a temporal resolution of 0.72 seconds. The preprocessing pipeline is described in [19] where the primary steps include motion correction, high-pass temporal filtering, and artefact removal [9].

### 3.2 Dense to Low-rank + Sparse connectomes (DLS)

As shown in Figure 1 and as discussed earlier, the strong similarity between the ICA maps and the whole-brain diffusion embedding maps suggests that both methods are presenting the same dominant global-linear patterns present in the DC, rather than the connectivity gradients intended to be uncovered by the embedding approach. To address this, and model the true gradients, in the DLS approach, we adopt a “low-rank + sparse” decomposition to separate global-linear low-rank components of the DC matrix indicative of functional segregation, from gradient-like connectivity profiles which should be both high-rank and sparse. We then apply local embedding techniques such as ISOMAP and UMAP, to the estimated sparse matrix, to find whole-brain connectivity embeddings, i.e., gradients.

Figure 2 shows an example DC that demonstrates how the components should appear after decomposition: the low-rank matrix **L** should contain block structures, whereas the sparse matrix **S** should resemble a full-rank matrix but with many zeros.

**Figure 2.**
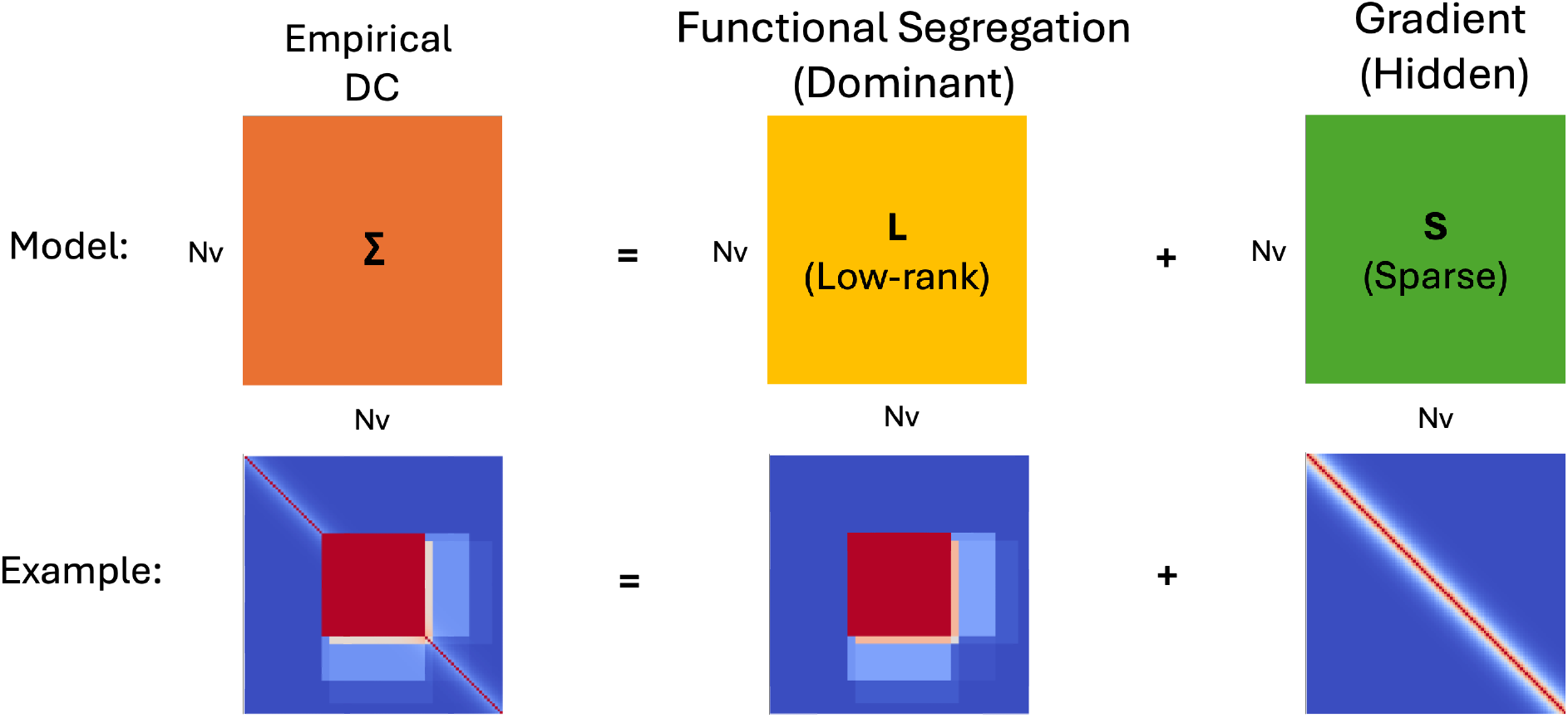
The empirical dense connectome can be decomposed into two components: one representing the global-linear connectivity profile that is low-rank, and another sparse matrix that captures local, continuous variations in connectivity where these variations are the primary focus of gradient modelling.

To estimate **L** and **S**, we solve the following optimisation, known as Principal Component Pursuit [3]:

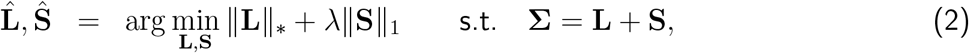

where ∥.∥_*_ is the nuclear norm of a matrix and ∥∥_1_ denotes the sum of the absolute values of the matrix entries. It has been shown in [3] that, under fairly broad conditions, the solutions to (2) with 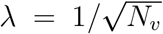 are guaranteed to converge to the true underlying low-rank and sparse matrices that constitute the empirical DC. These conditions boil down to (i) the low-rank connectome matrix **L** is not too sparse, and (ii) the (sparse) gradient connectivity matrix **S** is not low-rank. Optimisation can be done via alternating-directions using an augmented Lagrangian method on the dual form of the problem:

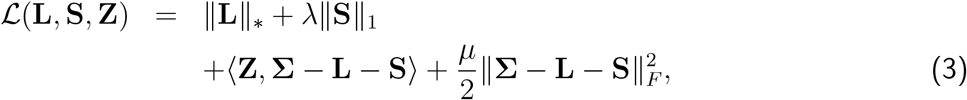

where ⟨., .⟩ is the matrix inner product and **Z** and *µ* are the Lagrange multiplier matrix and the penalty parameter, respectively, used to enforce the constraint **Σ** = **L** + **S. Algorithm** 1 summarises all the steps (including optional symmetrisation for faster convergence) taken to estimate the unknown matrices **L** and **S**. In this algorithm, D_1*/µ*_ is a singular value soft-thresholding function with threshold 1*/µ* and S_*λ/µ*_ is an element-wise soft-thresholding function with threshold *λ/µ*.

Following optimisation, the estimated matrix **S** is fed to nonlinear manifold learning methods, here either ISOMAP (global) or UMAP (local/global), to estimate the connectivity gradients. Although the algorithm is “parameter-free”, in the sense that *λ* has a theoretically prescribed value, deviations from the model assumption in real data could break this. Cross-validation can be used to optimise *λ*, but here we instead use cross-validation to choose the maximum possible rank of **L** (denoted by *r*_*m*_). A greedy search can be used in this case due to the discrete nature of this hyperparameter.

One final detail is that in order to perform this cross-validation, the model (“predicted”) DC must be formed. This involves transforming the embedding distances using the E2C transform (see 3.3) to match with the sparse component of the DC. We also add an additional global multiplicative factor (denoted by *w*^*^) before combining with the low-rank matrix to further improve the model DC. The full pipeline for gradient estimation and DC reconstruction in DLS is shown in Figure 3.

**Figure 3.**
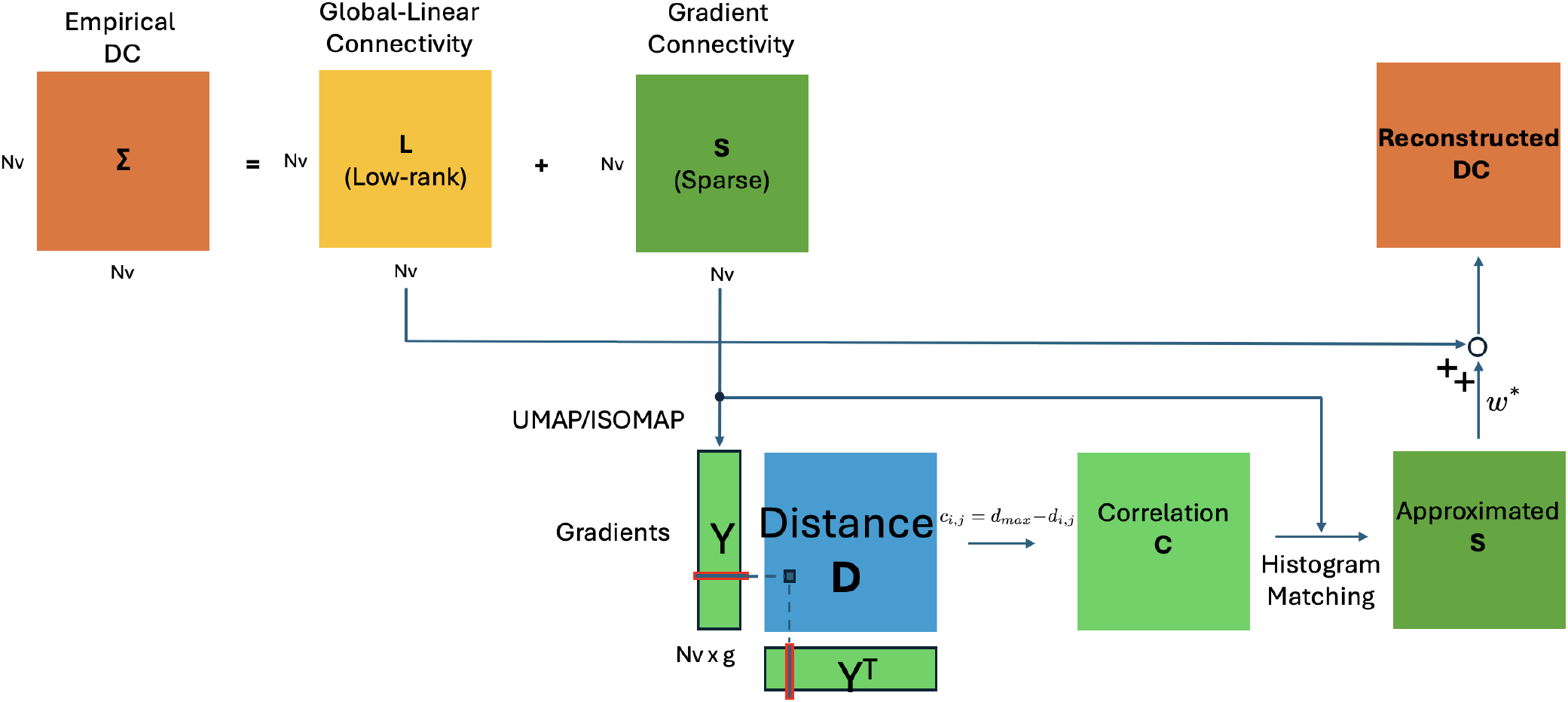
DLS Pipeline: The “L+S” decomposition is first applied to extract the sparse connectivity matrix that captures the fine-grained, continuous structure of the connectivity profile as well as the low-rank connectivity that represents the global functional segregation among brain ROIs. Subsequently, nonlinear embedding methods (such as ISOMAP or UMAP) are used to compute low-dimensional gradient maps 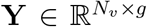. To evaluate how well the estimated gradients represent the original empirical DC, a global histogram matching-based E2C function is employed to approximate the sparse connectivity matrix from the gradients. This estimated gradient-based connectivity is then optimally combined with the low-rank matrix **L** to reconstruct the DC. Finally, the similarity between this reconstruction and either the input DC or a held-out test DC (assessed within a cross-validation framework) provides a quantitative measure of performance quality.

### 3.3 Embedding to Connectome (E2C)

To compare different methods for modelling the dense connectome, we will use a reconstruction loss. To do so, we will compare the empirical DC with the modelled DC using a correlation metric. However, as our approach is not a generative model, we first need to map the low-dimensional embedding back to a model dense connectome. To achieve this we use nonparametric global histogram matching, which we refer to as Embedding-to-Connectome (E2C). This is done by first transforming the distance matrix in the embedding space into a similarity matrix by replacing each pairwise distance *d*_*ij*_ with max_*ij*_(*d*_*ij*_) − *d*_*ij*_. Then nonlinear (but monotonic) global histogram matching is used to transform these similarities to match the distribution of the original 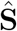 matrix.

## 4 Results

### 4.1 Synthetic Data

We generated a synthetic brain slice containing 4 square ROIs of equal size (25 × 25 voxels each). To construct ground truth gradients, and under the assumption that these ROIs are functionally related, we generated the same random time series (with *T* = 2, 000 time points) in all 4 regions, thus creating a one-to-one connectivity mapping. We also applied a spatial Gaussian smoothing kernel with *σ* = [*σ*_*x*_, *σ*_*y*_] = [1, 1] to these time series to introduce spatial autocorrelation. Hence, the correlation structure with reference to a seed voxel now changes smoothly across space, and underlying connectivity matrix is full rank with gradual transitions within the ROI rather than sharp boundaries. This procedure produces ground truth gradients within each ROI that take the form of two orthogonal (because of isotropic smoothing), smoothly increasing patterns, that are identical across ROIs (as the initialisations are identical). Because the smoothing is isotropic, rotated versions of these gradients are also valid solutions, since isotropic Gaussian smoothing acts equally in all spatial directions and therefore, preserves the same covariance structure under rotation. Additionally, to demonstrate the presence of global-linear patterns arising from functional segregation, we added a rank-2 data matrix to the gradient data plus relatively weak random noise. Each rank in the added matrix models the co-activation of two ROIs as indicated in the spatial maps in Figure 4 (A) with a shared time series. Figure 4 illustrates the generative process used to construct the dataset.

**Figure 4.**
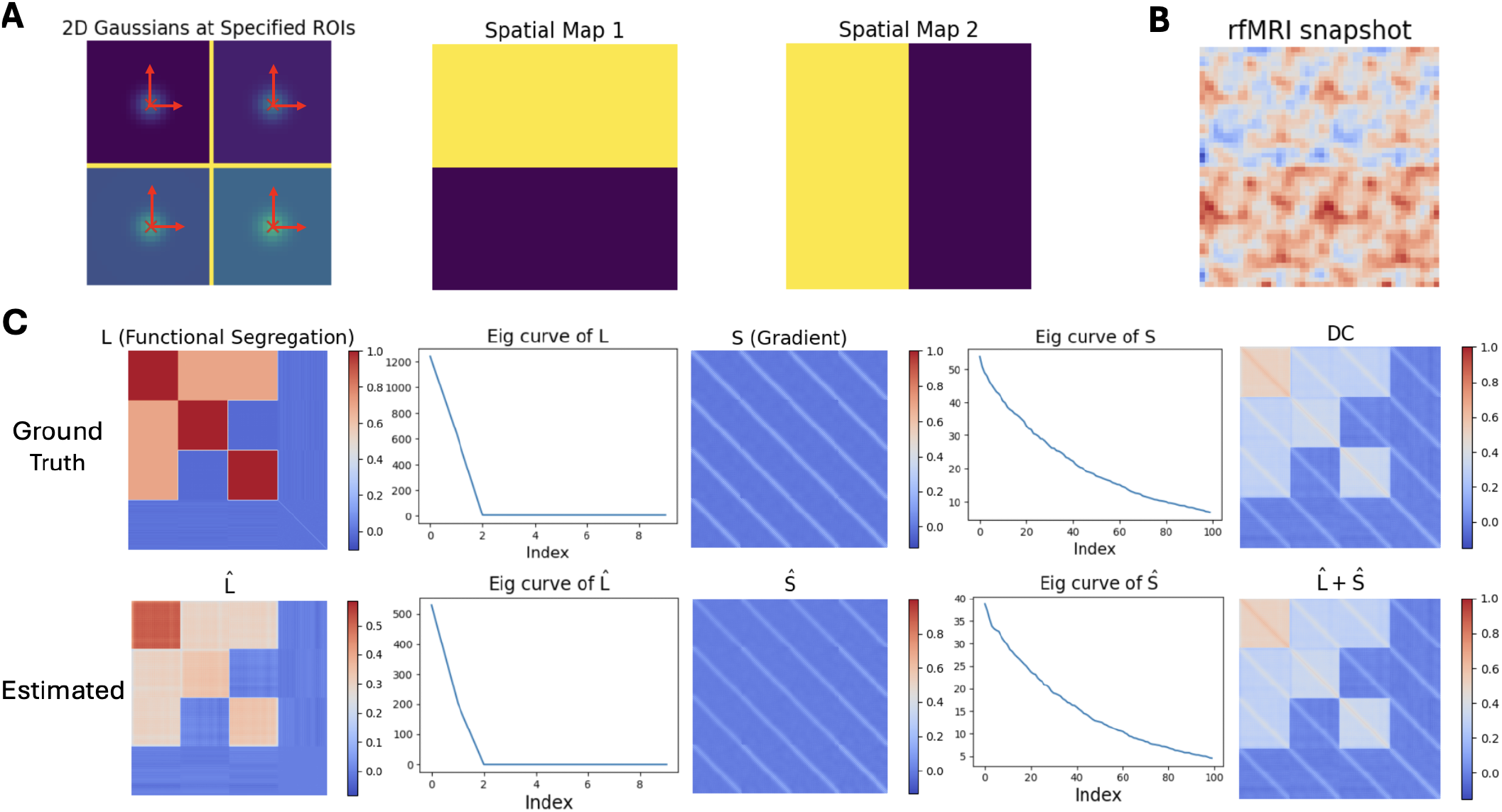
(A) From left to right: The simulated brain slice contains four ROIs. Isotropic Gaussian smoothing produces two-dimensional gradients within each ROI, which are identical across ROIs. Any rotated orthogonal directions are also valid solutions. The next two images show spatial maps in which each highlighted region (yellow vertices) shares a random time series, illustrating the effect of functional segregation. (B) A single snapshot of the generated data over time, visualised in the spatial domain. As shown, all ROIs remain synchronised in their spatial patterns, while the low-rank component introduces contrast variations. (C) The dense connectome for each component of the data (ground truth vs estimated). Functional segregation yields a rank-2 connectome, while the gradient produces a sparse connectome in which the eigenvalue spectrum contains many non-zero elements, trending toward a full-rank matrix. The combination of the two results in a dense connectome that exhibits both lowrank structure and gradient-rich characteristics. The estimated connectomes using the proposed DLS approach nicely resemble the ground truths.

For comparison, we contrast our results with a low-rank approximation method based on ICA and the global diffusion embedding method employed in [15] (hereafter referred to as the “Gradient Only” approach). As discussed earlier, the DLS approach can be optimised with respect to the *r*_*m*_ parameter that limits the rank of the matrix 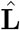. To estimate this parameter, we examine the value at which applying nonlinear embedding to the 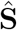 matrix, and its combination with 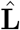, provides a better approximation of the DC compared to using only a low-rank approximation with the same number of components. We assume that any improvement should be detectable even with a single gradient component.

Figure 5 demonstrates that at *r*_*m*_ = 2, DLS with UMAP (with *g* = 2 gradients on top of *r*_*m*_ = 2) outperforms the purely low-rank approximation of the DC with the same number of dimensions (*r*_*m*_ = 4). This result confirms the presence of only two dominant global-linear patterns in the data, consistent with the process in the generative model. By fixing this parameter and applying DLS with UMAP/ISOMAP, we obtain the gradients shown in Figure 6, which can be compared with the whole-brain Gradient Only (third row) and ICA (last row). As shown in Figure 6 (A), the gradients estimated by DLS (particularly when using UMAP for nonlinear embedding) result in two orthogonal gradient directions (without this constraint being explicitly enforced in the loss function) that align closely with the ground truth gradients. Importantly, these gradients are consistent across all ROIs. Note that the orthogonality of the estimated gradients, if needed, can be imposed by certain types of nonlinear embedding approaches (in our case, ISOMAP and UMAP do not impose this constraint), and is not internally enforced by DLS. Looking at the scatter plots in Figure 6 (B) and visualising the two gradients against each other, we observe that DLS places related points from the four ROIs at the same locations while preserving the geometry of the ROI configuration in a rectangular slice. The colour map derived from the first gradient (PC1) further illustrates the expected linear increase in the embedding.

**Figure 5.**
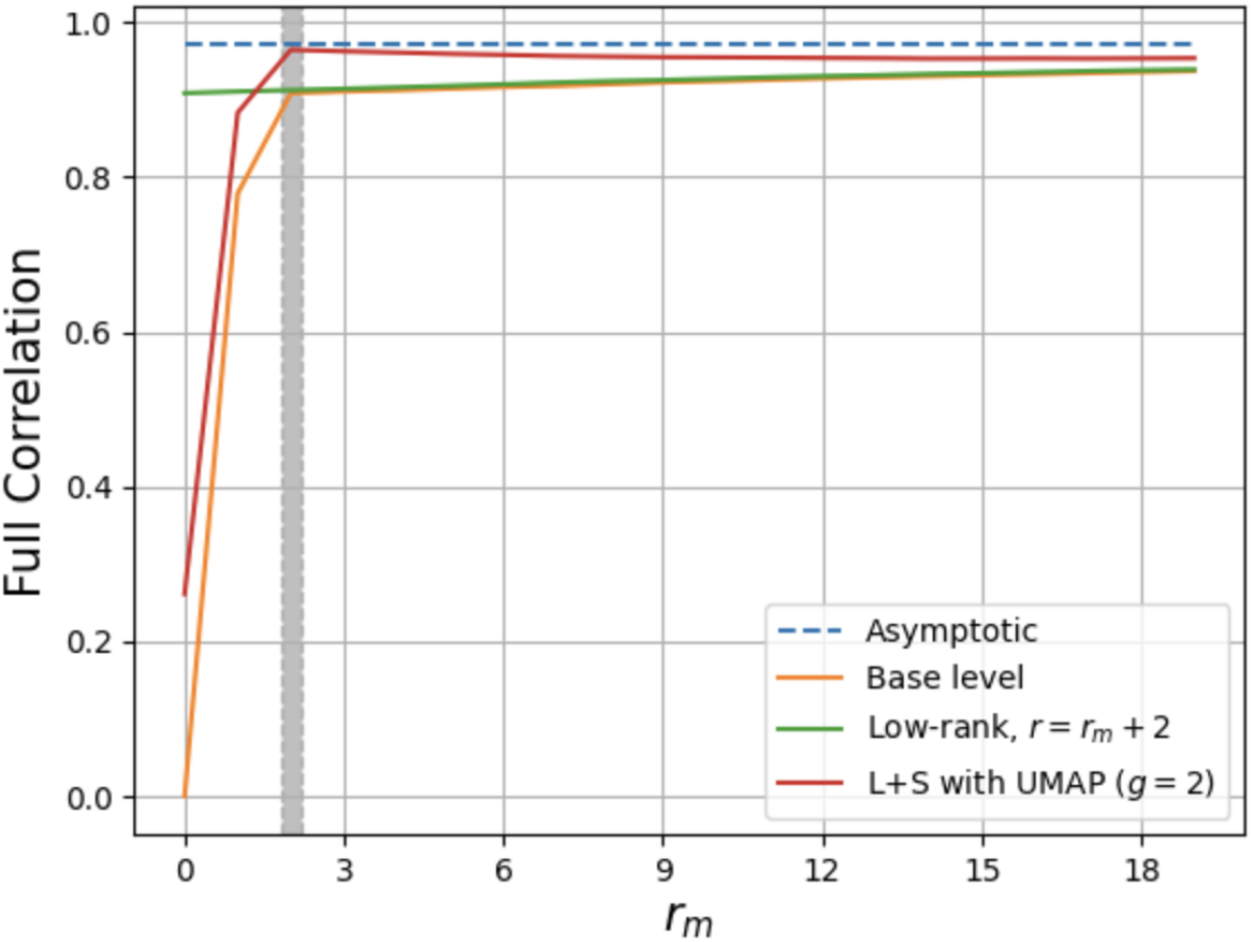
Optimising the maximum rank (*r*_*m*_) in 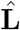 via cross-validation. The orange curve shows the correlation between 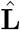 and empirical DC when the rank is limited by *r*_*m*_ (this is the base level as it just takes into account the low-rank estimate). The green curve shows the same metric when the maximum rank for 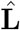 is increased to *r*_*m*_ + 2 (for the sake of a fair comparison with gradients). The red curve shows correlation of 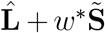 with empirical DC, where only 2 dimensions are used for the sparse matrix reconstruction 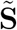 (with the same *r*_*m*_ constraint in 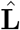). The best *r*_*m*_ where UMAP with gradient estimation outperforms the low-rank approximation in DC reconstruction with the same degree of freedom is found at *r*_*m*_ = 2 (shown in a gray vertical box). This is indeed the same number of spatial maps used in the generative process to creat global low-rank terms. Thus, these cross-validations help identify which parts of the DC are relevant to gradients and which suffice for modelling global-linear behaviour.

**Figure 6.**
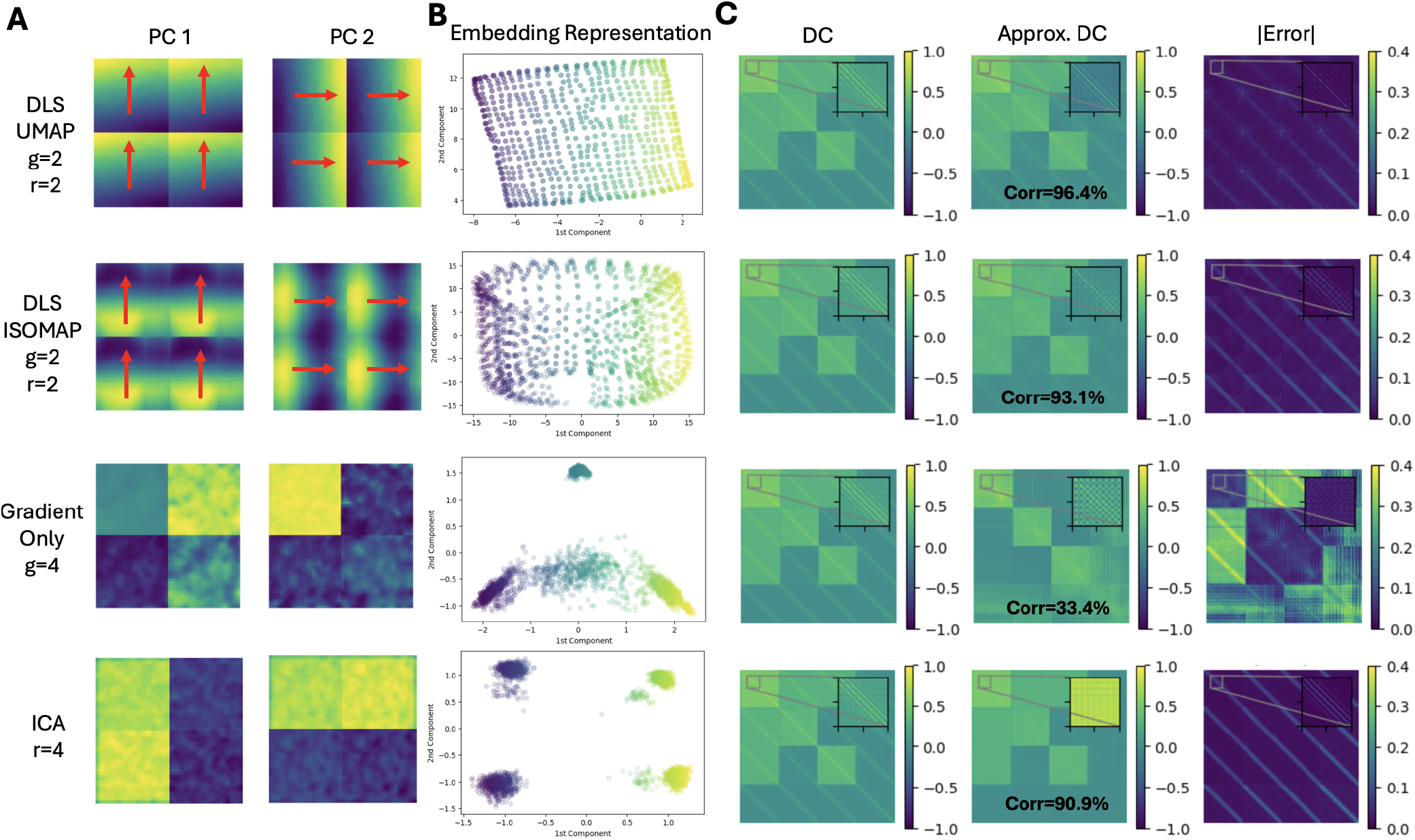
(A) The principal gradient maps and organisational directions in the four ROIs (shown as red arrows), are presented for the DLS variants, the whole-brain gradient only analysis [15], as well as the global-linear patterns, i.e., spatial ICA maps. (B) Scatter plots of the two gradients plotted against each other, colour-coded by PC1. Using DLS, the spatial geometry of each ROI is preserved in a rectangular configuration. The same points from different ROIs are mapped to the same locations, reflecting that their underlying gradients are identical. (C) Approximated DCs vs empirical DC, and their difference (Error), along with their calculated correlation (cross-validated).

Figure 6 (C) shows that DLS with UMAP achieves a strong recovery of the DC (Corr = 96.4%) using only two gradients and a low-rank term (r=2). DLS with ISOMAP also demonstrates nearly competitive performance (Corr = 93.1%), though with minor misalignment between ground truth and estimated gradients, likely due to its global behaviour. The error between the empirical DC and its approximation by DLS shows that the remaining minor error is mostly around modelling the gradient terms (diagonal elements).

As predicted, the results of the gradient only analysis is significantly influenced by global-linear terms in the data. The first two gradients primarily reflect the separation of ROIs from one another, without capturing any shared gradients across ROIs. This leads to irrelevant organisational directions and highlights only functional segregation. The ICA of the same data with 4-dimensional embedding (more visualisations in Figure S1) also separates the ROIs in almost the same pattern. This observation illustrates why the global gradient approach presented in [15] produces results similar to the ICA method, as shown in Figure 1. When applying the nonparametric E2C function to the results of these existing approaches, the DC reconstruction is poor (Corr = 33.4% and 90.9%) compared to the DLS. The error in DC reconstruction using ICA primarily arises from inaccuracies in the gradient terms. In the gradient-only approach, this error results from a combination of both global-linear and continuous variations. The superior performance of DLS arises from its ability to effectively and simultaneously model both the global-linear and the gradient-rich (and potentially sparse) terms in the empirical data, as illustrated in Figure 4 (C).

### 4.2 HCP Data

A random subset of 100 HCP subjects was selected, and we used their minimally preprocessed restingstate fMRI data in our analysis. For this study, subcortical areas were excluded. To form the empirical DC, we used Pearson correlation for the sampled version of brain voxels/vertices (randomly 25% of the voxels/vertices were selected) to reduce computational complexity. Note that the full connectome, without any sampling strategy, can also be fed into the DLS pipeline; however, this would be more demanding in terms of computational resources. We divided the subjects into two equal groups (50/50 subjects) to perform cross-validation and identify the optimal hyperparameters, such as the *r*_*m*_ parameter, within the DLS framework. We hypothesised that, at the optimal *r*_*m*_, incorporating two gradient components will yield better performance than adding two further low-rank components. This approach follows the same logic applied to the simulated data. The choice of two global gradients is motivated by previous findings from task-fMRI analyses, which shows that brain ROIs typically exhibit one or two dominant gradients [10]. As illustrated in Figure 7, incorporating two gradients produced a clear performance peak at *r*_*m*_ = 9, which corresponds to the upper bound of the rank in 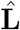. Applying DLS to the empirical DC in the training set returns the estimated **L** and **S** matrices. It is not surprising that the low-rank component shows a strong correlation with the empirical DC (probabilistic scatter plot in Figure 8). This finding once again conceptually demonstrates that the influence of global-linear patterns in the DC at the whole-brain scale should be treated with caution; otherwise, it may result in misleading gradient maps. Figure S2 presents the empirical DC and its eigen-curve in comparison with the estimated **L** and **S** matrices. As expected, both matrices comply with the assumptions underlying their estimation process—namely, **L** exhibits a low-rank structure, whereas **S** is sparse (and not low-rank).

**Figure 7.**
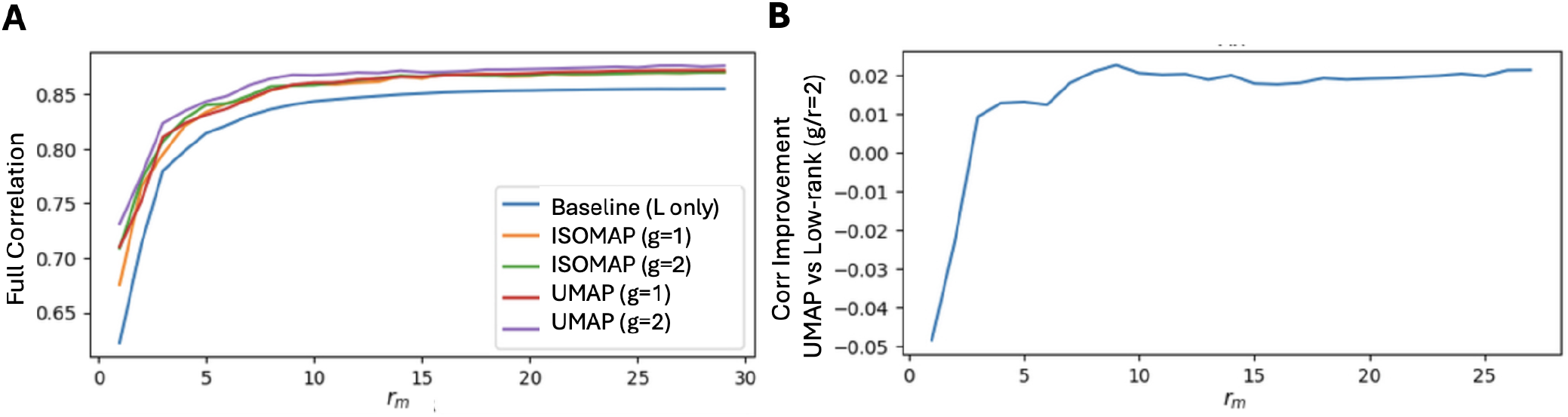
Cross-validation results for optimising *r*_*m*_ in the HCP dataset. (A) Cross-validated correlation between the reconstructed DC and the empirical DC as a function of *r*_*m*_. DLS with UMAP (*g* = 2) yields the best performance. (B) At *r*_*m*_ = 9, the improvement in DC reconstruction using DLS with UMAP (*g* = 2) reaches its peak when compared with a purely low-rank approximation incorporating the same number of components.

**Figure 8.**
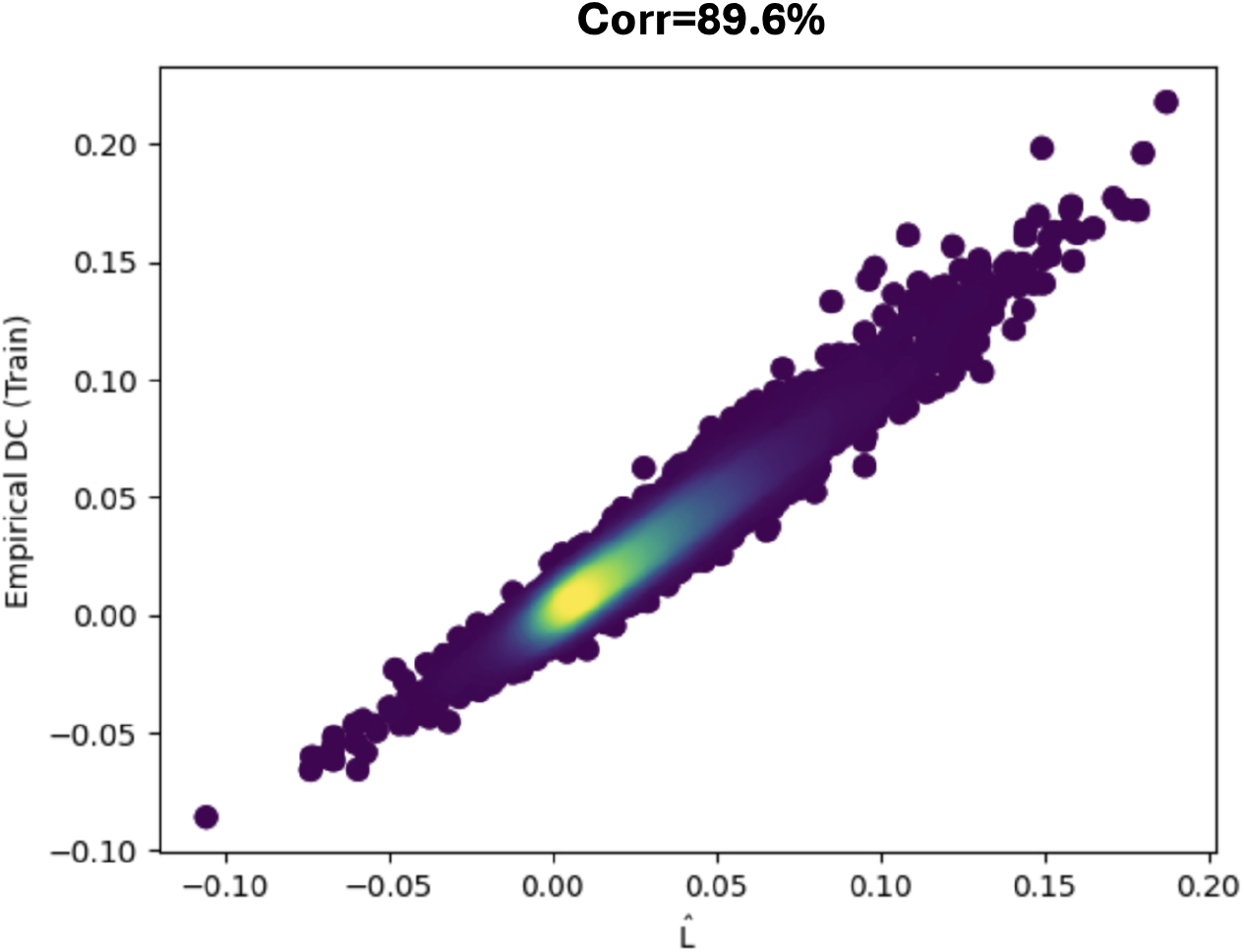
The estimated **L** with *r*_*m*_ = 9 from the HCP dataset shows a strong correlation with the empirical DC. The probabilistic scatter plot of sampled entries from both matrices indicates that the majority of values can be well approximated by the low-rank component of the DLS, while the gradient is embedded within the residual part of the DC which is presented in the 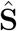.

Figure 9 on the other hand, shows that for the majority of brain regions, global-linear modelling is sufficient to explain their connectivity profiles, as a large portion of the maps in B is colour-coded in blue (smaller values) while most areas in A are colour-coded in white-red (large values). As shown in Figure 9 (B) in red (larger values), only a subset of regions requires more sophisticated embedding approaches beyond low-rank approximation, such as gradient modelling, to characterise their connectivity patterns (this aligns with the sparsity constraint). Those areas include visual, motor, and sensory areas, as well as some regions within the default mode network.

**Figure 9.**
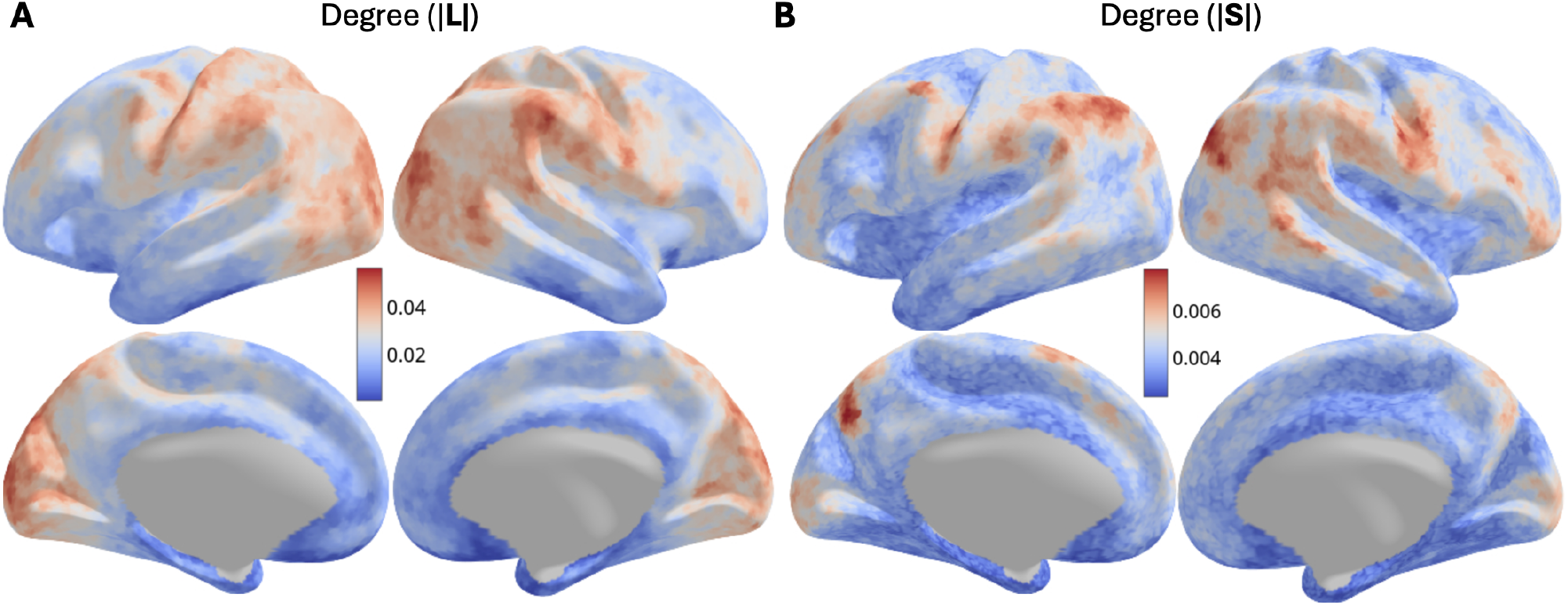
Applying DLS to the empirical DC from the HCP dataset yields two matrices: the low-rank matrix 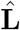 and the sparse matrix 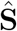. (A) The average absolute connectivity strength map, known as the average degree map (all positive values), in 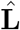, and (B) the average degree map in 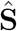. These maps illustrate which regions of the brain require gradient modelling, in addition to the low-rank approximation, to better capture the connectivity profile in a hybrid manner.

To demonstrate the importance of disentangling functional segregation from gradient-rich information in the DC, we examined the posterior–anterior direction in V1 (Figure 10 and more results later in Figure 13). Given the known visual eccentricity gradient present along this axis, substantial changes in the connectivity profile were expected. This change is clearly visible in the sparse connectivity profiles for three seed location choices, which primarily capture local and strong connections. In contrast, the corresponding connectivity profiles in 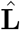 show minimal variability. This again explains why underlying gradients are not easily identifiable in whole-brain gradient analysis approaches that directly operate on DC embeddings. We next investigate whether gradient embeddings derived using the DLS framework (capturing true local variations) reveal meaningful large-scale organisation when projected onto the cortical surface, in a manner analogous to the visualisation presented in [15]. To do so, 3 distant points in the embedding space were selected as reference points and assigned distinct colours: Red, Green, and Blue. The central point of the spectrum was also highlighted in yellow as the new reference point, then the remaining points in proximity to these references were colour-coded according to their relative distances (Figure 11). This visualisation reveals partially (the difference will be discussed later) the same conclusion made in the gradient only result, where the primary sensory motor areas, i.e., the primary visual (V1), primary motor (M1), primary somatosensory (S1), and primary auditory (A1) regions occupy all the extreme ends of the spectrum from the gradient perspective developed in this paper. These regions correspond to those previously identified as requiring gradient analysis in conjunction with their global-linear patterns to fully characterise the connectivity profiles they exhibit (Figure 9). The top two gradients of the cortical areas, extracted by DLS using both ISOMAP and UMAP in the embedding step, are shown in Figure 12. To enable a meaningful comparison between the two embeddings (ISOMAP vs UMAP), we aligned the UMAP embedding to that obtained from ISOMAP through rotation and translation within the embedding space. It should be noted that such a transformation does not affect the UMAP loss function, as the relative positions of all samples are preserved and the resulting embeddings are also valid solutions. Examining Gradient 1 and Gradient 2 (without implying any specific order), both DLS variants display largely consistent gradients for the DC, with minor differences attributable to the local versus global perspectives inherent in the respective embedding approaches. One immediate observation is that these maps differ from those produced by ICA or gradient only approaches. The question then arises as to how these maps can be validated? One method of validation is to examine the derived maps within regions of interest (ROIs) for which the expected gradient patterns are already known. Examples of such ROIs include the primary motor cortex (M1), where a top–down gradient is expected (known as the somatotopic map), and the primary visual cortex (V1), where the eccentricity gradient is anticipated along the posterior–anterior axis [10]. To do so, we mask out all vertices outside these ROIs from the gradients visualised in Figure 12 (one at a time for each ROI), and then rescale the colour map for the remaining embeddings to better represent the variations within each ROI. Figure 13 confirms that, when these global gradients are masked to M1 and V1 regions, they exhibit similar profiles as somatotopic and eccentricity maps, whereas in the gradient only method this consistency was not observed. As mentioned before, the orthogonality of the estimated gradients are not enforced by ISOMAP or UMAP. One may wish to examine the results with this additional constraint on gradients. Hence, results for DLS with Spectral Embedding (SE) [2] were also provided to show the results are consistent across well-known nonlinear embedding techniques. While this visualisation qualitatively validates the gradients estimated by DLS, we may still wish to quantitatively validate these gradients. Another way of validation is to examine the capability of these estimated gradients in approximating the DC, as they play a collaborative role with low-rank components, i.e., through 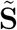: the output of global histogram matching, in 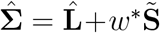. Given this formulation, it is evident that in the DLS framework, both the global-linear low-rank components (Figure S3) and the gradients are necessary for the connectome reconstruction.

**Figure 10.**
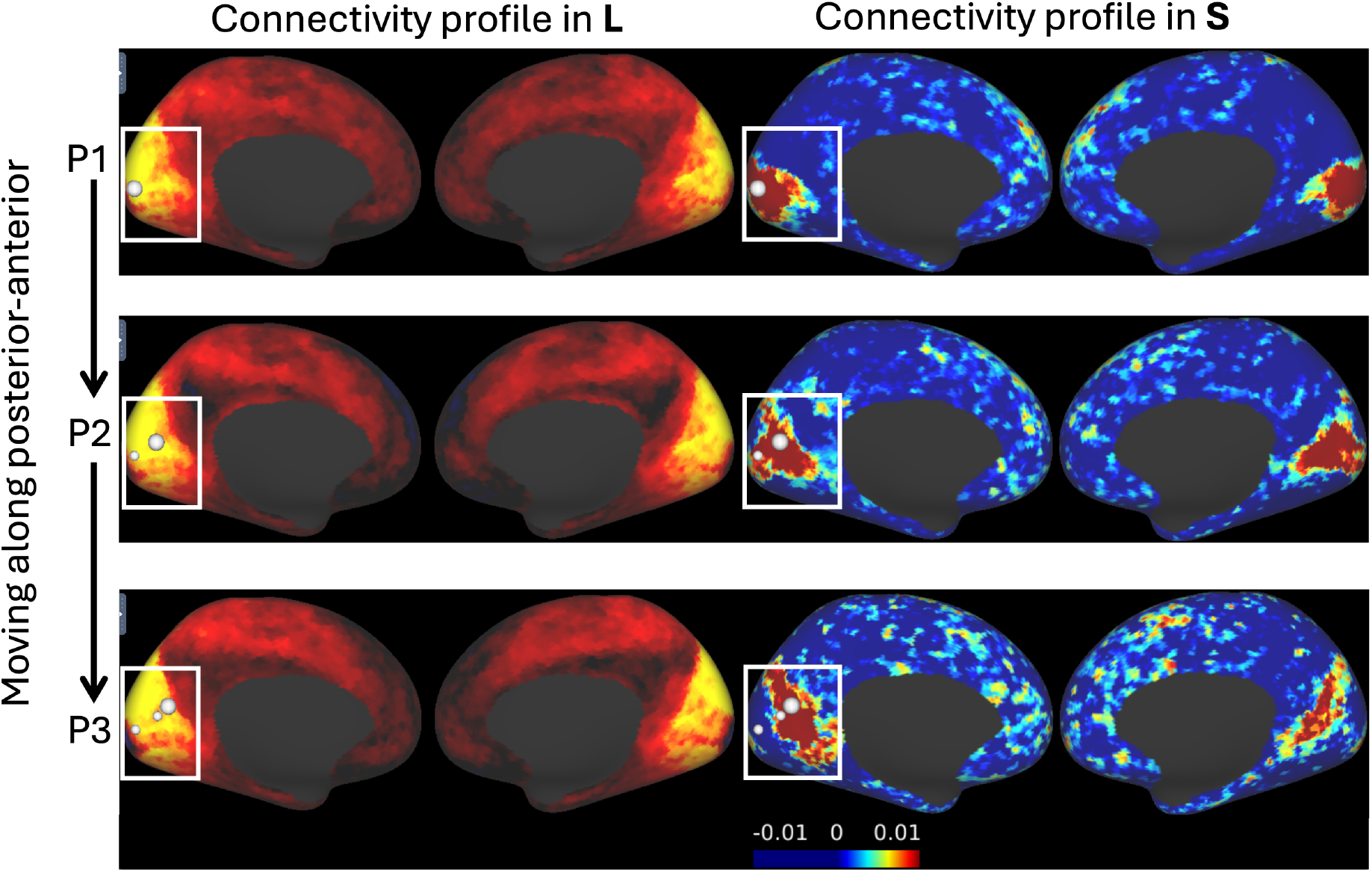
An example of differing behaviours in connectivity profile changes between estimated **L** and **S** ( HCP dataset) is illustrated. Moving from P1 to P3 via P2 along the posterior–anterior axis in the V1 area does not produce any change in the connectivity profile within 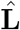, but reveals a local variation in the connectivity pattern represented by 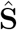 Given the knowledge about the presence of the eccentricity gradient along this axis, working on 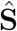 results in more meaningful gradients.

**Figure 11.**
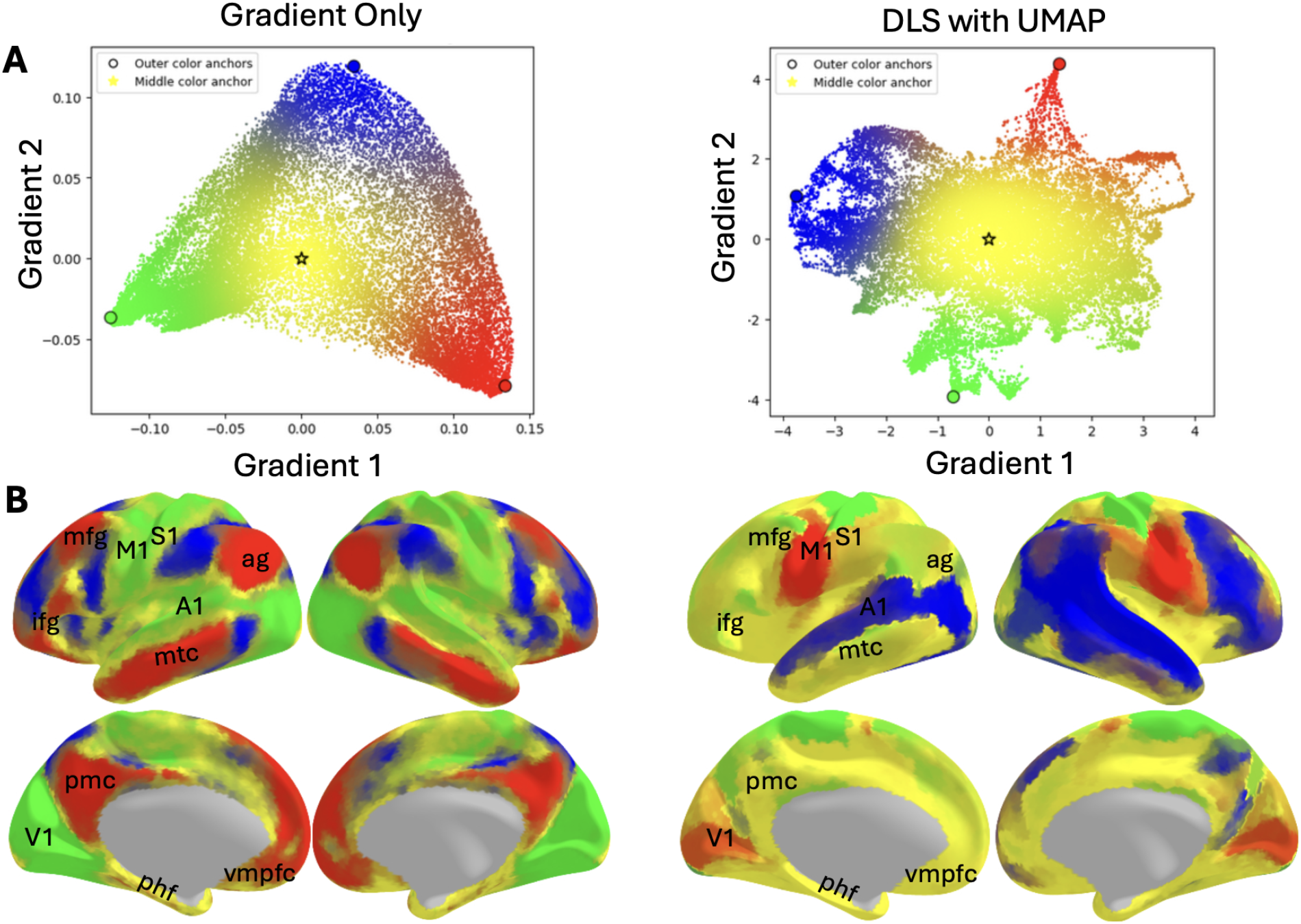
Scatter plots of the HCP-related gradients estimated by gradient only approach and DLS and their clustering/variation behaviours on the brain surface. Colours from the scatter plots in (A) are presented on the cortical surface for anatomical orientation in (B). While the gradient-only approach (with normalised angle kernel) identifies three clusters with only a few vertices in the intermediate space of the embedding (yellow), DLS spans the entire spectrum more uniformly, representing a continuous variation across the brain, consistent with gradient definition. The yellow colour in DLS (B) indicates regions situated near the centre of the spectrum, without a clear membership to any of the three clusters formed at the outer ranges. A1, primary auditory; ag, angular gyrus; ifg, inferior frontal gyrus; M1, primary motor; mfg, middle frontal gyrus; mtc, middle temporal cortex; phf, para-hippocampal formation; pmc, posteromedial cortex; S1, primary somatosensory; V1, primary visual; vmpfc, ventromedial prefrontal cortex.

**Figure 12.**
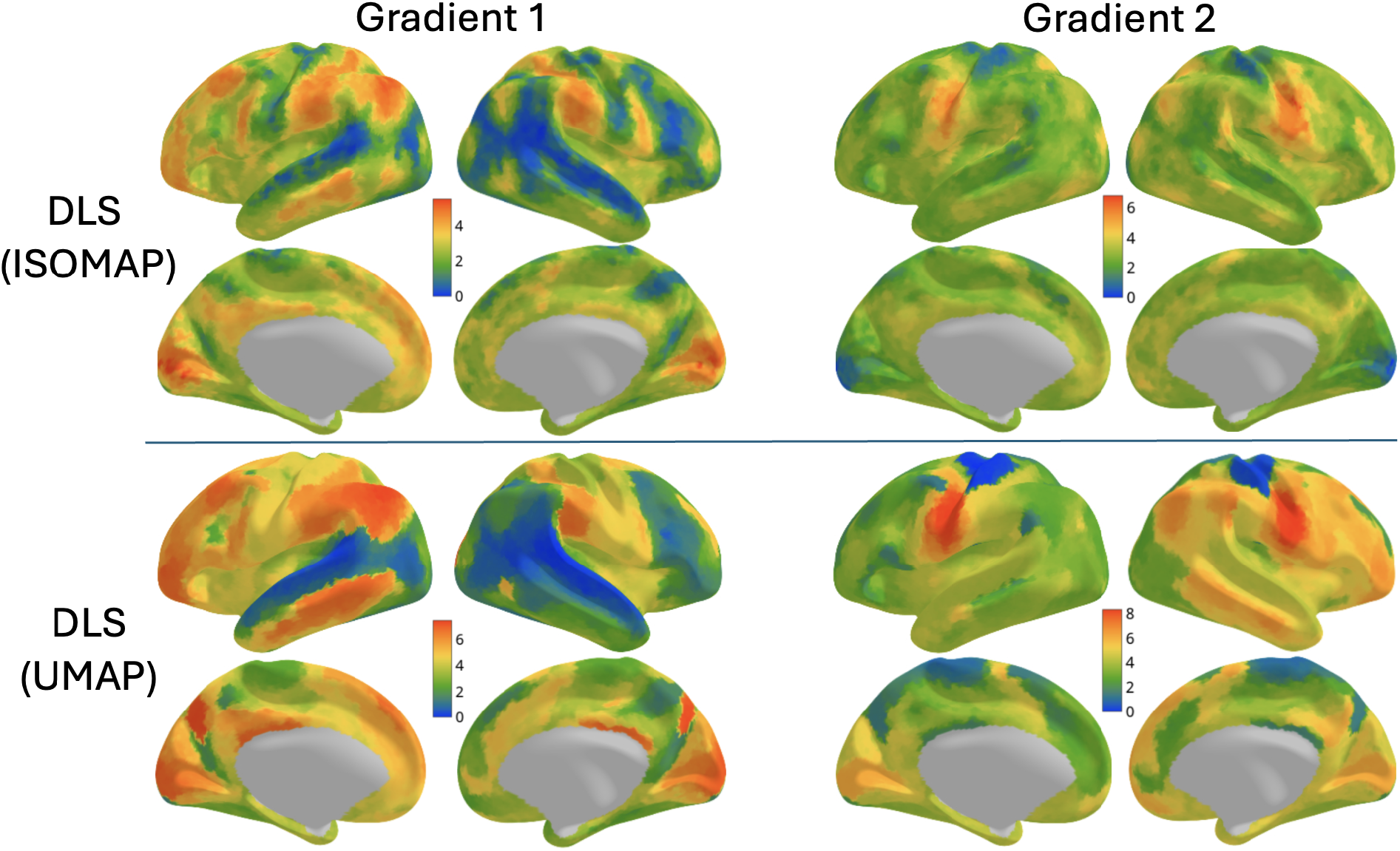
Global Gradients estimated by DLS variants from the HCP dataset. The results are consistent with one another, and both estimated global gradients exhibit the same major patterns of variation across the brain, e.g., top-down variation in Gradient 2 within M1 and S1 areas.

**Figure 13.**
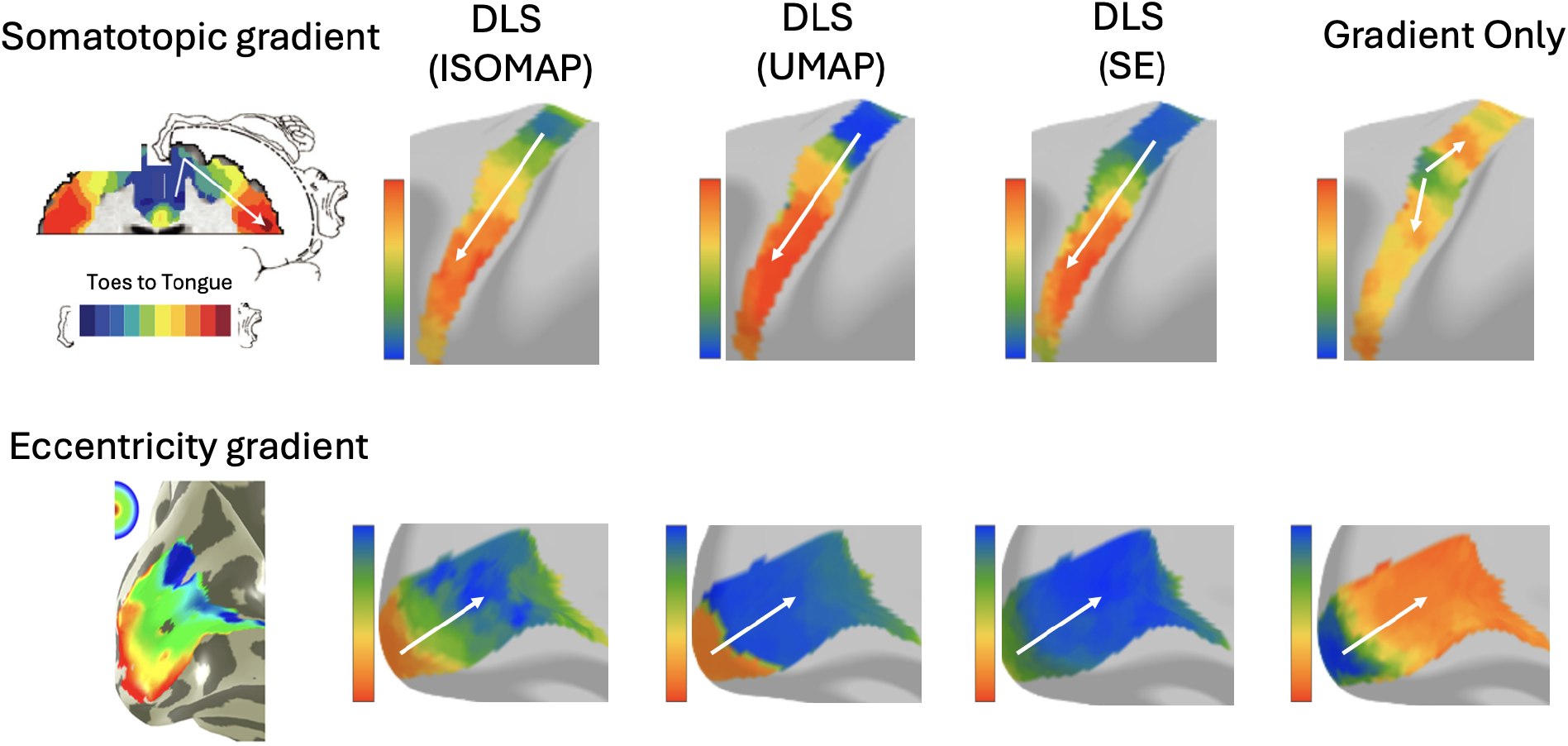
Local gradients are obtained by masking the global gradients for a target ROI. The gradients derived from the DLS variants resemble the known gradient patterns in the M1 and V1 regions. Somatotopic gradient map of M1 (top) and retinotopic V1 eccentricity map (bottom) as the dominant gradients are from [27] and [10], respectively.

Table 1 shows that, among all DC modelling approaches—including the low-rank approximation and gradient only methods—the DLS approach, which is a hybrid method, achieves the highest DC recovery rate, with UMAP yielding 86.8 % correlation with the validation DC using only nine global-linear patterns combined with two global gradients. While the correlation improvement of DLS over the gradient-only approach is substantial (46.0% → 86.8%), the improvement over the low-rank approximation is more moderate (84.5% → 86.8%). This is because the first nine low-rank components dominate the DC, and all competition between the global and local approaches occurs in the residual modelling, which accounts for only a limited portion of the DC approximation. If the objective were to model the residual part 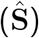 rather than the DC itself, this difference would be more apparent in the correlation metric (Table 2).

**Table 1.** Cross-validated correlation value between the empirical DC and the approximated DC in HCP dataset. *k*: total number of dimensions, *r*: number of low-rank dimensions, *r*_*m*_: the cap on the number of low-rank dimensions, *g*: number of gradients.

| Method/# Components | Condition | $k = 9$ | $k = 11$ |
| --- | --- | --- | --- |
| Gradient Only ([15]) | $r = 0$ | 46.2% | 46.0% |
| Low-rank Approx. | $g = 0$ | 84.1% | 84.5% |
| DLS (ISOMAP) | $r_m = 9$ | 84.1% | 85.8% |
| DLS (UMAP) | $r_m = 9$ | 84.1% | **86.8%** |

**Table 2.** Correlation value between the 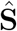 and its approximation using different methods in HCP dataset.

| Method | Low-rank Approx. | DLS (ISOMAP) | DLS (UMAP) |
| --- | --- | --- | --- |
| **Correlation with $\hat{S}$** | 24.1% | 44.9% | **54.0%** |

Figure 14 shows all the approximated DCs displayed next to each other, where the DLS approach stands out in terms of its similarity to the empirical DC and its ability to capture the fine details of the DC.

**Figure 14.**
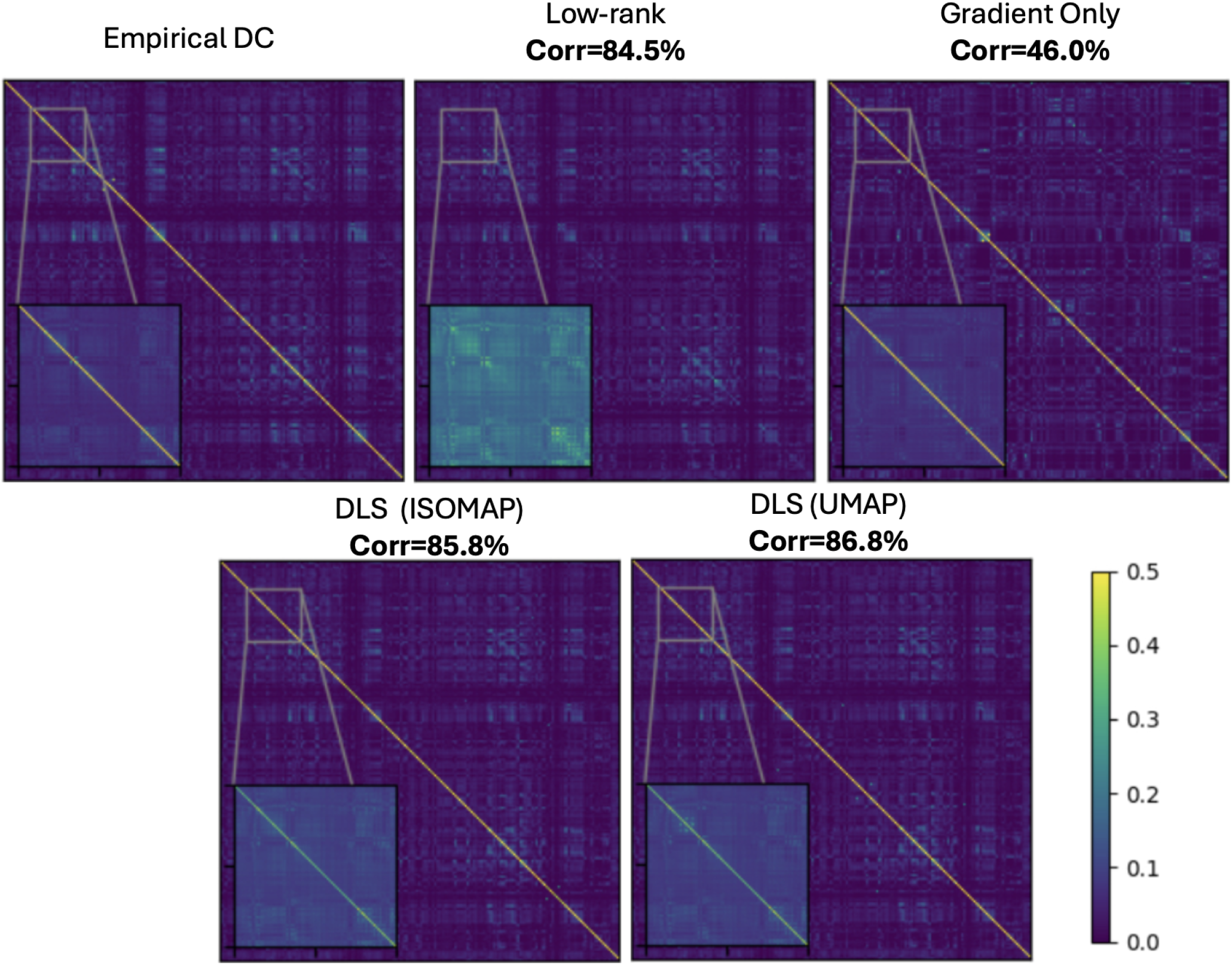
Approximated DCs vs empirical DC in HCP dataset with k=11. The upper range of the colormap was clipped to enhance contrast.

## 5 Discussion

A common approach in DC modelling is the treatment of brain data using either global-linear approaches (e.g., ICA) or local embedding methods (e.g., gradient methods). However, neither of these approaches can fully explain the DC in practice. The former requires a large number of components to fully describe the data accurately. On the other hand, the latter fails to account for functional segregation, which is the dominant part of the data, and hence does not reveal brain gradients that explain the actual neural organisation well. Figure 13, for example, illustrates such a failure in the M1 area due to its small number of vertices and the influence of other similar regions. Here, we developed a hybrid approach, termed DLS, which respects both global-linear and local-nonlinear patterns in the data. Given the data structure, the method identifies which portions should be described using linear versus local nonlinear models. By reconciling these two traditionally separate approaches, we estimated a global brain organisation that, from a gradient perspective, resembles known local neural organisation in a continuous form. As a beneficial side effect, the DC can be explained with a smaller number of parameters, as each parameter is assigned to model the aspects of the data it best represents. For example, using data from 50 HCP subjects, we found that 9 global-linear components and 2 local gradients (capturing the heterogeneity in the data) achieved 86.8% similarity with the left-out DC, with the remaining dissimilarity largely reflecting noise variations between the training and test sets. Our analysis shows that for achieving this level of DC reconstruction, if we use standard approaches in isolation as typically applied in the literature, more than 500 ICA components or global gradients (∼50x) are needed.

In Figure 9, the data visualisation shows that while some areas can be fully explained by global-linear patterns, some are more complex than being modelled by such abstract models. Here is where hybrid approaches such as DLS boosts the modelling performance, as it adopts both global-linear and local-nonlinear approaches to explain the data. Our further investigation shows that, although the low-rank component contributes equally to both inter- and intra-hemispheric connectivity profiles modelling, the average gradient connectivity within each hemisphere is primarily driven by inter-hemispheric connections (Figures S4, S5). This observation is consistent with the notion that gradient information is largely local and reflects relationships among neighbouring vertices, whereas the global-linear terms capture large-scale organisation (top level representation) irrespective of local variations.

Given that in the literature and Figure 11 (A) it was shown that the principal gradients derived from the gradient only approach expand from regions serving primary sensory motor functions to the other regions, including transmodal regions [11], our results also show almost the same organisation with slightly different configuration. The embedding visualisation of the gradient only technique often exhibits a clustering behaviour of the connectome rather than a continuous variation along the embedding dimensions. Figure 11 (A) shows that the embedding in the gradient only configuration places almost all vertices within the ROIs adjacent to each other (forming a cluster) instead of distributing them along a continuous spectrum. In contrast, our results in Figure 11 show that DLS spans the entire range, with most vertices uniformly located in the spectrum. This demonstrates that the maps are continuous, with less clustering effect often caused by global-linear low-rank patterns in the data. Moreover, in contrast to the findings presented in [15], the areas associated with cognition and higher-level functions, including transmodal regions, are predominantly located in the central part of the spectrum. Note that while this seems inconsistent with known hierarchical order, i.e., the brain’s structure and function are organised in levels, where lower levels process simple or local information, and higher levels integrate, abstract, or control that information, transmodal areas here may act as hubs to functionally connect regions at the extremes of the spectrum, i.e., primary sensorimotor areas. On the other hand, the low-rank representation of the data helps to clarify the hierarchical organisation, as transmodal regions that in human are known as Default Mode Network (DMN) are immediately highlighted as a network distinct from the others.

Now that DLS brain gradients have been illustrated for two well-known ROIs and for the whole-brain, our framework provides the tools needed to explore gradients in territories that have so far remained relatively uncharted with respect to gradient structure, e.g., hippocampus [6]. This opens the door to examining what these gradients represent within their respective ROIs. As illustrated in Figure 13, DLS successfully recovered the eccentricity gradient (an established feature of early visual cortex) demonstrating its sensitivity to biologically meaningful organisation. Beyond these familiar patterns, the approach should expose additional, as yet unexplained gradients, which can now be interrogated through targeted task-based fMRI experiments to uncover their functional significance.

Functional brain gradients are emerging as sensitive biomarkers for early-stage neurodegenerative diseases [7]. They have the potential to capture subtle disturbances in neural networks that arise before clear clinical symptoms emerge, offering insight into underlying disease mechanisms and their progression. In Alzheimer’s Disease (AD), for example, alterations in functional gradients within the medial parietal cortex have been linked to CerebroSpinal Fluid (CSF) markers such as phosphorylated tau and total tau, as well as cognitive decline in asymptomatic individuals at increased risk of AD [24]. Similarly, in Parkinson’s Disease (PD), reorganisation of functional gradients in the basal ganglia has been associated with emerging motor deficits and cognitive alterations [4]. Collectively, these findings emphasise the potential of functional gradients as Imaging-Derived Phenotypes (IDPs) for detecting early neural dysfunction, even before pronounced structural changes are evident. As such, they provide a promising foundation for developing biomarkers that support earlier diagnosis, inform prognosis, and potentially guide therapeutic strategies in neurodegenerative diseases.

Although the DLS framework demonstrates promise in capturing local and global neural organisation and delineating the full spectrum of brain regions, it relies on a hyperparameter that requires optimisation through an exhaustive search, which can be computationally demanding, though in an integer space. To expedite this process, we employed a sampling strategy that substantially reduces matrix dimensionality. While the L+S decomposition is computationally efficient, the subsequent nonlinear embedding stage remains resource-intensive. Another consideration is that DLS is entirely data-driven and does not employ any parametric modelling, which may not be favourable in direct applicability in clinical contexts. The framework produces multiple brain maps which, if one wishes to approximate using spatial parametric models for their own application, must be fitted post hoc (potentially introducing error). Consequently, DLS currently lacks a generative model that would allow for estimation within a Bayesian framework. Furthermore, as DLS derives global gradients from spatially local connections, distant interactions only contribute when the nonlinear embedding is configured with sufficiently high number of neighbours. A mechanistic extension that more explicitly incorporates long-range connectivity would therefore be beneficial, given that brain gradients often reflect topographical organisation where a voxel/vertex is linked not only to its immediate neighbours but also to distant regions. Finally, the analysis presented in this paper relied solely on rfMRI data. It would be worthwhile to extend the method to multiple types of brain imaging data in order to achieve a multimodal whole-brain gradient analysis [25].

## Data and Code Availability

The data used in this manuscript come from the Human Connectome Project dataset: https://www.humanconnectome.org/. Python code for processing real data and generating simulations will be openly available on GitHub: https://github.com/arekavandi/DLS.

## Author Contributions

A.M.R.: Conceptualisation, Methodology, Validation, Formal Analysis, Writing—Original Draft, Visualisation. S.J.: Conceptualisation, Methodology, Supervision, Writing—Reviewing & Editing, Funding Acquisition. S.M.S.: Conceptualisation, Methodology, Supervision, Writing—Reviewing & Editing, Funding Acquisition.

## Acknowledgements

Data were provided by the Human Connectome Project, WU-Minn Consortium (Principal Investigators: David Van Essen and Kamil Ugurbil; 1U54MH091657) funded by the 16 NIH Institutes and Centers that support the NIH Blueprint for Neuroscience Research; and by the McDonnell Center for Systems Neuroscience at Washington University.

## Use of Generative AI

The manuscript was edited with the assistance of ChatGPT (OpenAI) to improve clarity and language.

## Supplementary Material

The supplementary material is included in this PDF file after References.

## A Appendix

**Figure S1.**
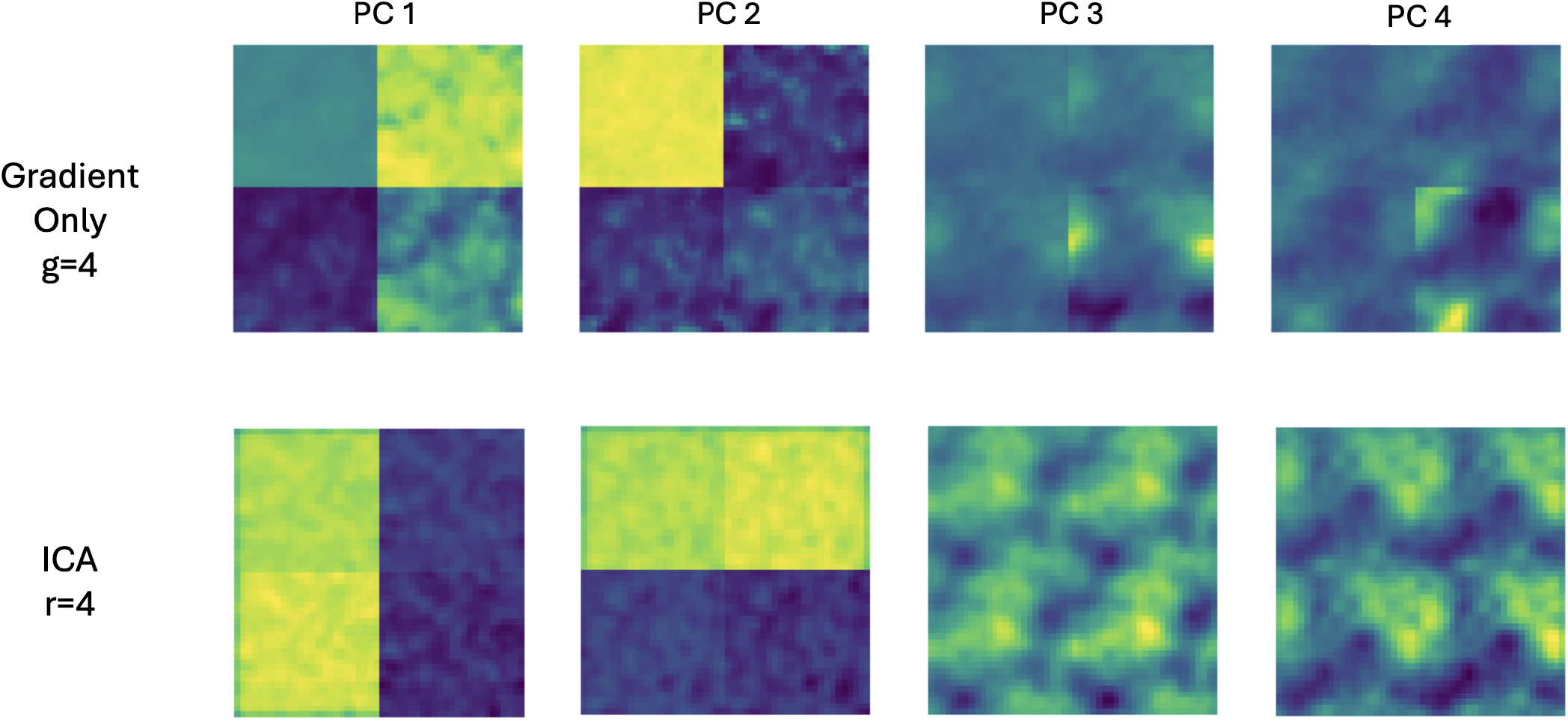
The top 4 principal gradients [15], as well as the spatial ICA maps for synthetic data.

**Figure S2.**
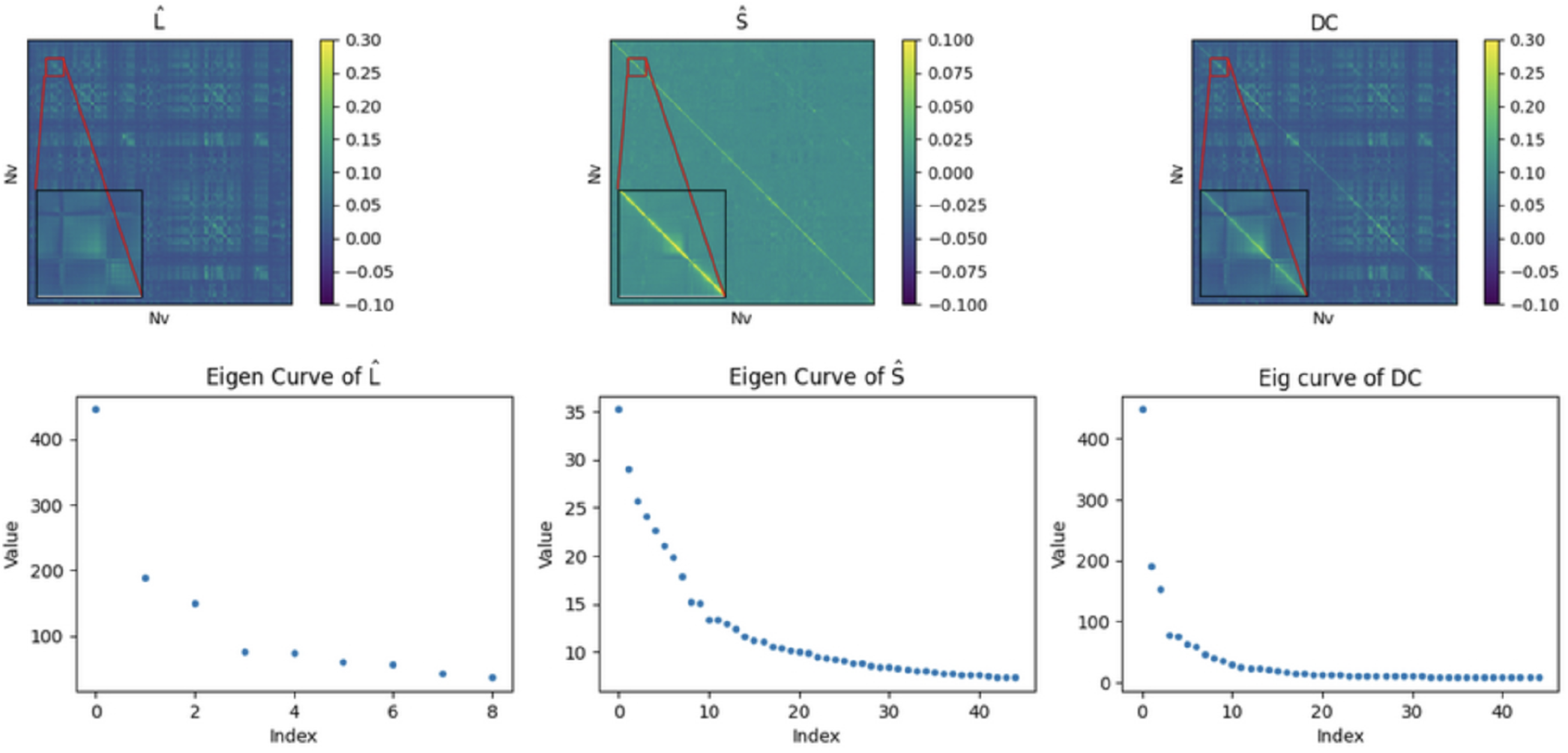
Estimated **L** and **S** matrices from HCP dataset, visualised together along with the empirical DC and their eigen-curves.

**Figure S3.**
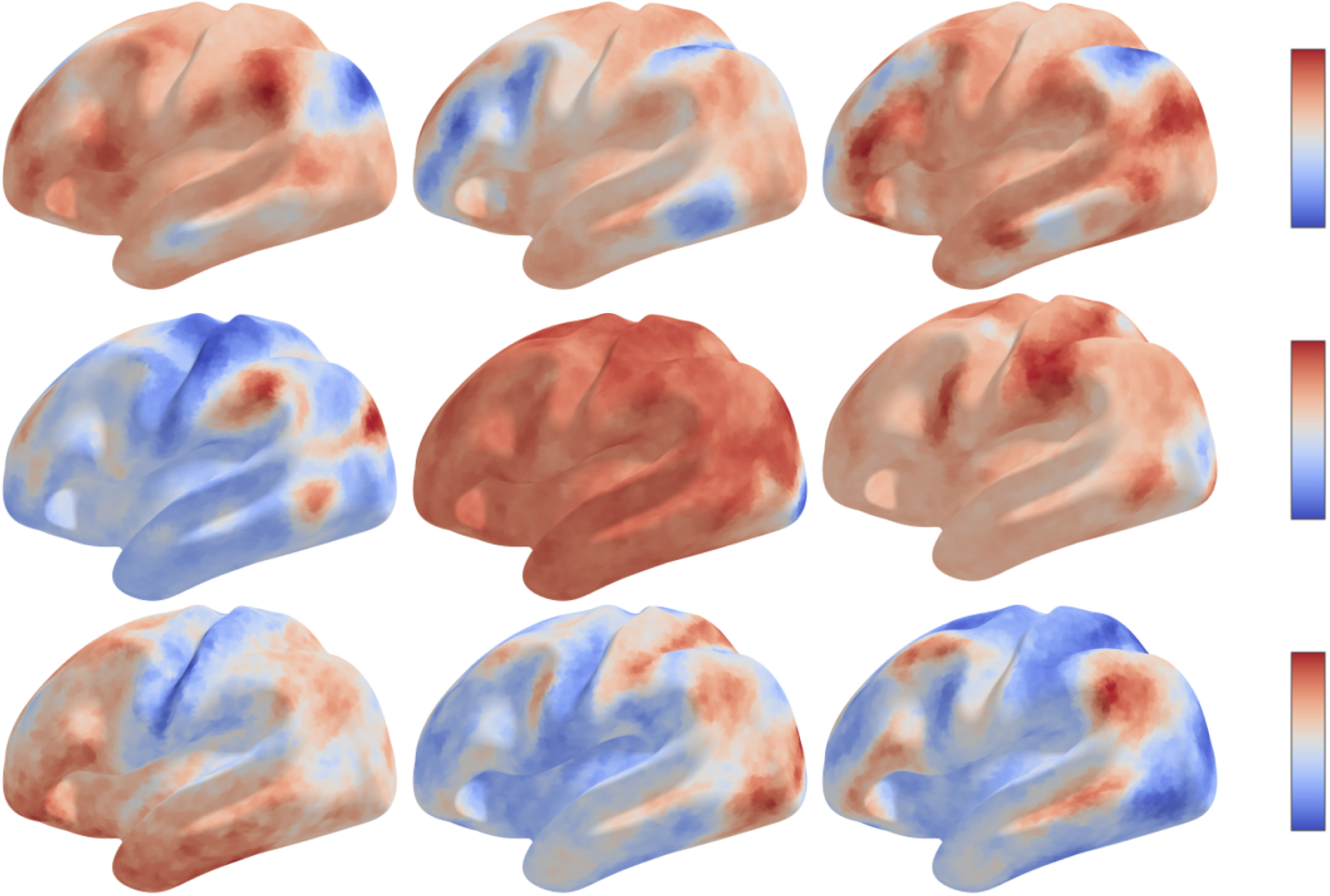
ICA components of 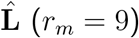 for HCP dataset.

**Figure S4.**
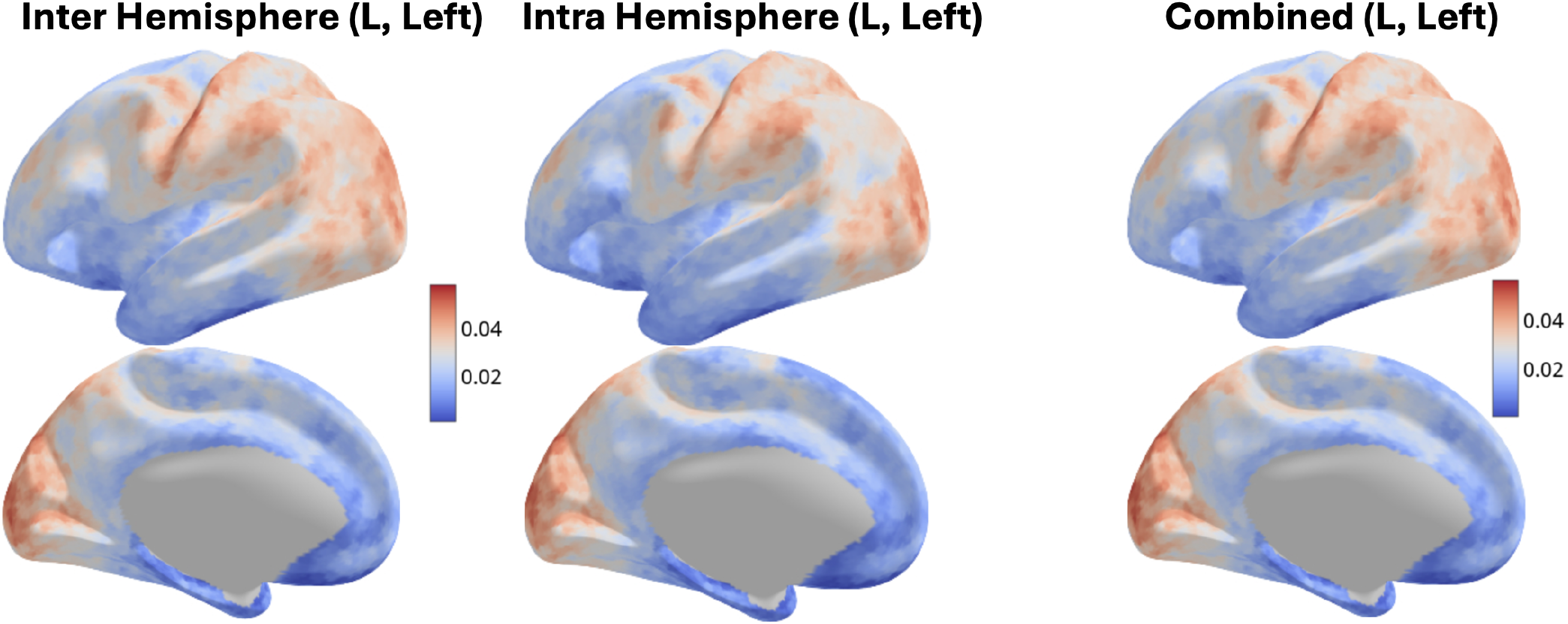
The average intervs intra-hemispheric vs combined connectivity maps in 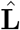 visualised for left hemisphere (HCP dataset).

**Figure S5.**
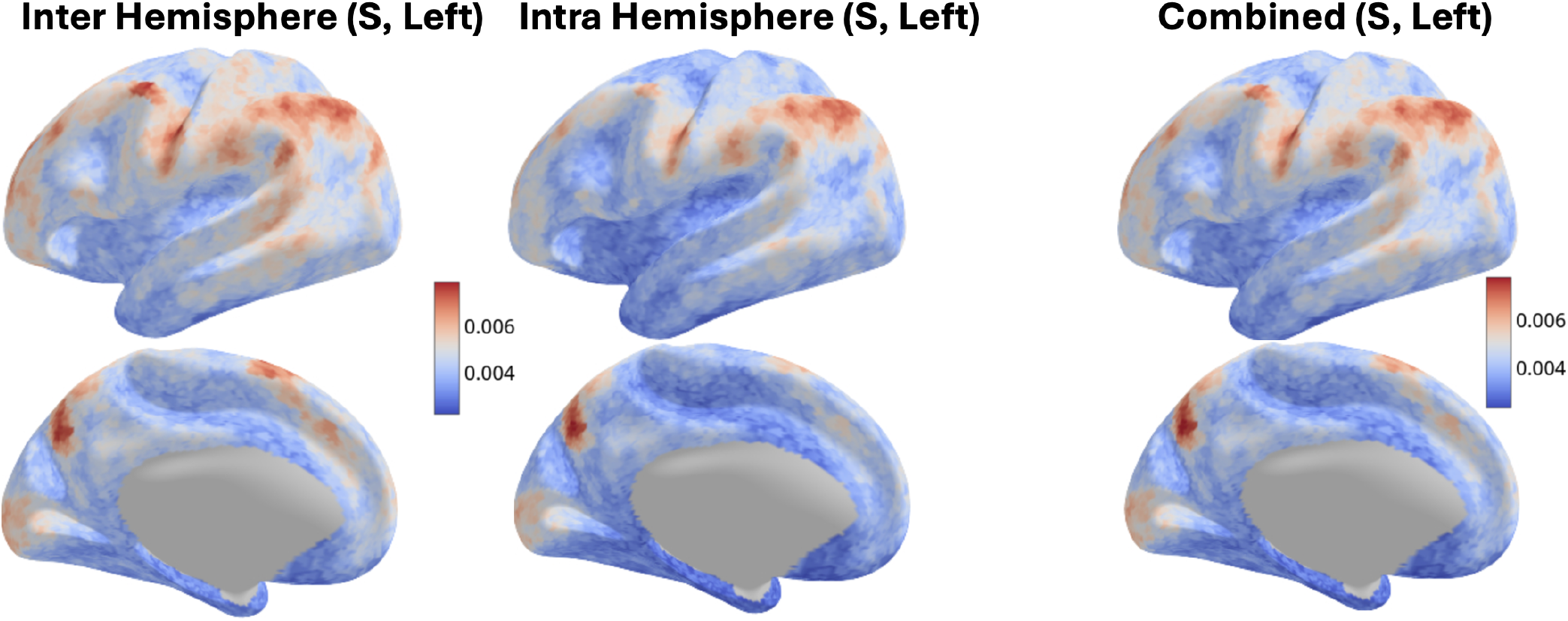
The average intervs intra-hemispheric vs combined connectivity maps in 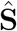 visualised for left hemisphere (HCP dataset).

